# Nodal secures pluripotency upon embryonic stem cell progression from the ground state

**DOI:** 10.1101/093880

**Authors:** Carla Mulas, Tüzer Kalkan, Austin Smith

## Abstract

Naïve mouse embryonic stem (ES) cells can readily acquire specific fates, but the cellular and molecular processes that enable lineage specification are poorly characterised. Here we investigated progression from the ES cell ground state in adherent culture. We utilised down-regulation of *Rex1::GFPd2* to track loss of ES cell identity. We found that cells that have newly down-regulated this reporter have acquired competence for germline induction. They can also be efficiently specified for different somatic lineages, responding more rapidly than naïve cells to inductive cues. Nodal is a candidate autocrine regulator of pluripotency. Abrogation of Nodal signalling did not substantially alter kinetics of exit from the ES cell state, but accelerated subsequent adoption of neural fate at the expense of other lineages. This effect was evident if Nodal was inhibited prior to extinction of ES cell identity. We suggest that Nodal is pivotal for non-neural competence in cells departing naïve pluripotency.

## INTRODUCTION

Pluripotency denotes a flexible cellular potential to differentiate into all lineages of the developing embryo. This property emerges in the epiblast of the pre-implantation blastocyst (Boroviak et al., 2014; Gardner, 1975; Rossant, 1975). After implantation, epiblast cells remain pluripotent while undergoing profound cellular and molecular changes in preparation for gastrulation (Smith, *in press*). In mice the post-implantation epiblast develops into a cup-shaped epithelium, the egg cylinder. Signalling cues from extra-embryonic tissues then pattern the egg cylinder to establish anterior-posterior and proximal-distal axes prior to lineage specification (Arnold and Robertson, 2009; Beddington and Robertson, 1998; Peng et al., 2016; Rossant and Tam, 2009; Thomas and Beddington, 1996).

In mouse the naive phase of pluripotency can be captured in culture in the form of embryonic stem (ES) cells (reviewed by Nichols and Smith, 2012). Dual inhibition (2i) of Mek1/2 and GSK3, in optional combination with the cytokine Leukemia Inhibitory Factor (LIF), allows mouse ES cells to maintain the transcription profile, DNA methylation status and developmental potential characteristic of the pre-implantation epiblast from which they are derived (Boroviak et al., 2014; 2015; Habibi et al., 2013; Leitch et al., 2013; Ying et al., 2008). ES cells in 2i are stable and relatively homogeneous, a condition referred to as “ground state” (Marks et al., 2012; Wray et al., 2010). Such uniformity in defined conditions provides an experimental system to characterise cellular and molecular events that generate multiple lineage-committed states from a developmental blank canvas.

ES cell progression from the ground state is initiated simply by removal of the inhibitors. In adherent culture this results predominantly in neural specification (Ying et al., 2003) or in a mixture of neural and mesoendodermal fates, depending on cell density (Kalkan et al., 2016). Previous studies have identified expression of Rex1 (gene name *Zfp42*) as a marker of undifferentiated ES cells (Betschinger et al., 2013; Kalkan and Smith, 2014; Leeb et al., 2014; Toyooka et al., 2008; Wray et al., 2010; 2011; Yang et al., 2012). In this study, we exploit a *Rex1*::*GFPd2* (*RGd2*) reporter cell line (Kalkan et al., 2016) to isolate cells at initial stages of progression from naïve pluripotency following release from 2i in adherent serum-free culture. We examine whether cells exiting the ES cell state guided by autocrine cues commit preferentially to a neural fate or exhibit competence for multilineage differentiation.

## RESULTS

### Multi-lineage differentiation capacity is retained after loss of naïve ES cell identity

In *Rex1::GFPd2* (RGd2) reporter ES cells, a short half-life GFP is expressed from the endogenous Rex1 (*Zfp42*) locus (Marks et al., 2012; Wray et al., 2011). Loss of the reporter coincides with downregulation of naïve pluripotency factors and functionally with extinction of clonal self-renewal capacity (Kalkan et al., 2016) (Figure S1A-D). GFP downregulation is asynchronous across the population. For at least 16 hours cell remain uniformly GFP positive (Kalkan et al., 2016). By 24hrs, however, GFP is expressed at variable levels and in a minority of cells the reporter is no longer detectable. These Rex1-negative cells have lost the capacity to resume self-renewal in 2i/LIF, whereas cells with high GFP retain comparable colony forming efficiency to cells taken directly from 2i (Figure S1C). We focussed attention on the character of cells 24hrs after 2i withdrawal, the first time point at which it is practical to isolate a substantial population of Rex1-negative cells by flow cytometry (Kalkan et al, 2016).

We first investigated capacity to form primordial germ cell-like cells (PGCLC). Previous studies have shown that undifferentiated ES cells are not directly competent for germline specification but must first transition to a transient epiblast-like (EpiLC) population which can then be induced to form PGCLC (Hayashi et al., 2011; Nakaki et al., 2013). The EpiLC population is obtained by transfer from 2i/LIF to N2B27 medium supplemented with ActivinA, bFGF and the serum substitute KSR for 48hrs (Hayashi et al., 2011). We assessed whether the first cells that exit the ground state in N2B27 alone exhibit competence to form PGCLC. For this purpose we used RGd2 ES cells transfected with a doxycycline (Dox)-regulatable expression construct containing the three germ line determination factors *Prdm1* (Blimp1), *Prdm14* and *Tfap2c* (Magnúsdóttir et al., 2012; Nakaki et al., 2013). Stable transfectants were withdrawn from 2i for 24hrs and the high and low GFP fractions isolated by fluorescence-activated cell sorting (FACS) (Figure 1A). For each fraction, 3000 cells were aggregated in non-adherent 96 well plates in medium containing 15% KSR with or without Dox (Nakaki et al., 2013). After 4 days, few cells co-expressing Blimp1 with Oct4 were present in aggregates from either population without Dox. Dox treatment did not increase the frequency of co-expression from Rex1-positive cells, but induced many double positive cells from the Rex1-negative fraction (Figure 1B and C). Dual expression of Blimp1 and Oct4 is a combination unique to PGCs and PGCLCs (Hayashi et al., 2011; Kurimoto et al., 2008; Nakaki et al., 2013). Furthermore, undifferentiated ES cells do not tolerate appreciable levels of Blimp1 protein (Magnúsdóttir et al., 2013). Quantitative image analysis confirmed more intense Blimp1 staining in cultures derived from Rex1-negative cells (Figure 1D). By RT-qPCR analysis we detected upregulated expression of endogenous *Prdm1* (Blimp1), along with *Prdm14, Tfap2c, Nanos3* and *Stella,* as well as maintenance of *Pou5f1* (Oct4) (Figure 1E). *T* (Bra) was induced transiently on day 2 as previously described for PGCLC induction (Figure 1E) (Nakaki et al., 2013). Thus ES cells that have newly exited the ground state under autocrine stimulation in defined conditions acquire competence for germline specification.

**Figure 1.**
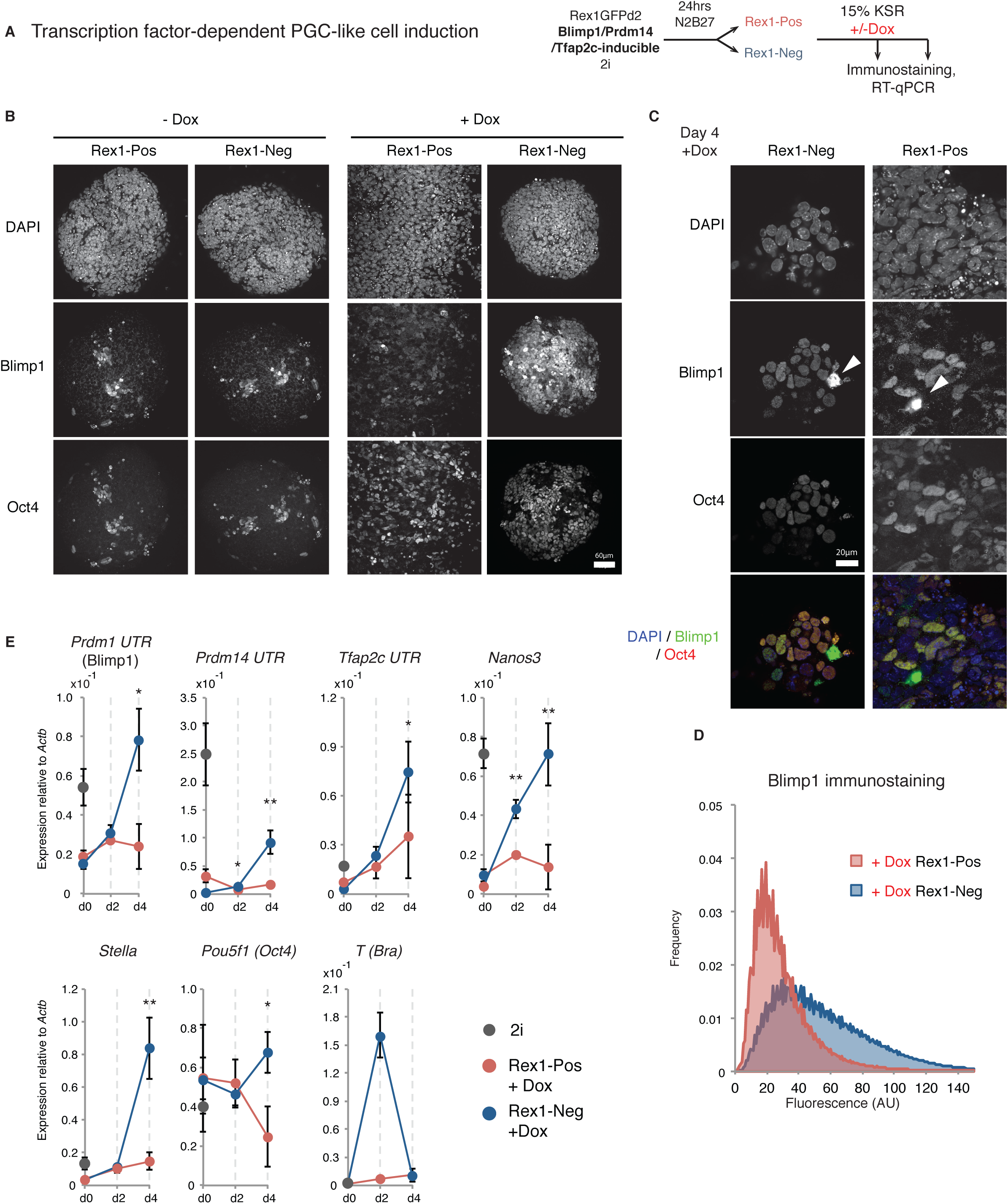
Acquisition of PGC-LC differentiation capacity. (**A**) Experimental set up for transcription-factor dependent PGC-like cell specification. (**B**) Expression of Blimp1 and Oct4 in day 4 aggregates differentiated in the presence or absence of Dox to induce transcription factor overexpression. Scale bar: 60μm (**C**) Zoom in of the expression of Blimp1 and Oct4 in day 4 aggregates differentiated in the presence or absence of Dox to induce transcription factor overexpression. Arrow heads show overexpression artefacts. Scale bar: 20μm (**D**) Quantification of the Blimp1 staining on day 4 in aggregates after addition of Dox. (**E**) RT-qPCR of endogenous PGC-associated transcripts. Mean and SD for 2 independent experiments shown, *p<0.01, **p<0.001 (Student t test). See also Figure S1

We then examined somatic lineage potential of Rex1-negative cells. Sorted fractions were plated in media that favour mesoderm, definitive endoderm or neural lineages respectively and the timing and efficiency of differentiation quantified.

ActivinA combined with GSK3 inhibition (GSK3(i)), elicits the upregulation of primitive-streak markers such as T (Tbra) in differentiating ES cells (Gadue et al., 2006; Tsakiridis et al., 2014; Turner et al., 2014)Morrison et al., 2015). We modified *RGd2* cells to express an mKO2 fluorescent reporter from the *T(Bra)* locus (Figure 2A). *T::mKO2* was not expressed in undifferentiated ES cells in 2i (Figure S2A), and not detected until day 3 of treatment with Activin plus GSK3(i). In contrast, Rex1-negative cells replated in the presence of ActivinA and GSK3(i) upregulated *T::mKO2* after one day and all cells were positive by day 2. Rex1-positive cells upregulated *T::mKO2* at an intermediate rate and some cells remained GFP-positive even after 3 days, indicating they remained undifferentiated and unresponsive to differentiation cues (Figure 2B). To test further mesoderm differentiation, we plated the sorted fractions in conditions that promote lateral mesoderm (Nishikawa et al., 1998; Yamashita et al., 2000). All populations gave rise to Flk1 positive/E-cadherin negative cells after 4-5 days (Figure 2C).

**Figure 2.**
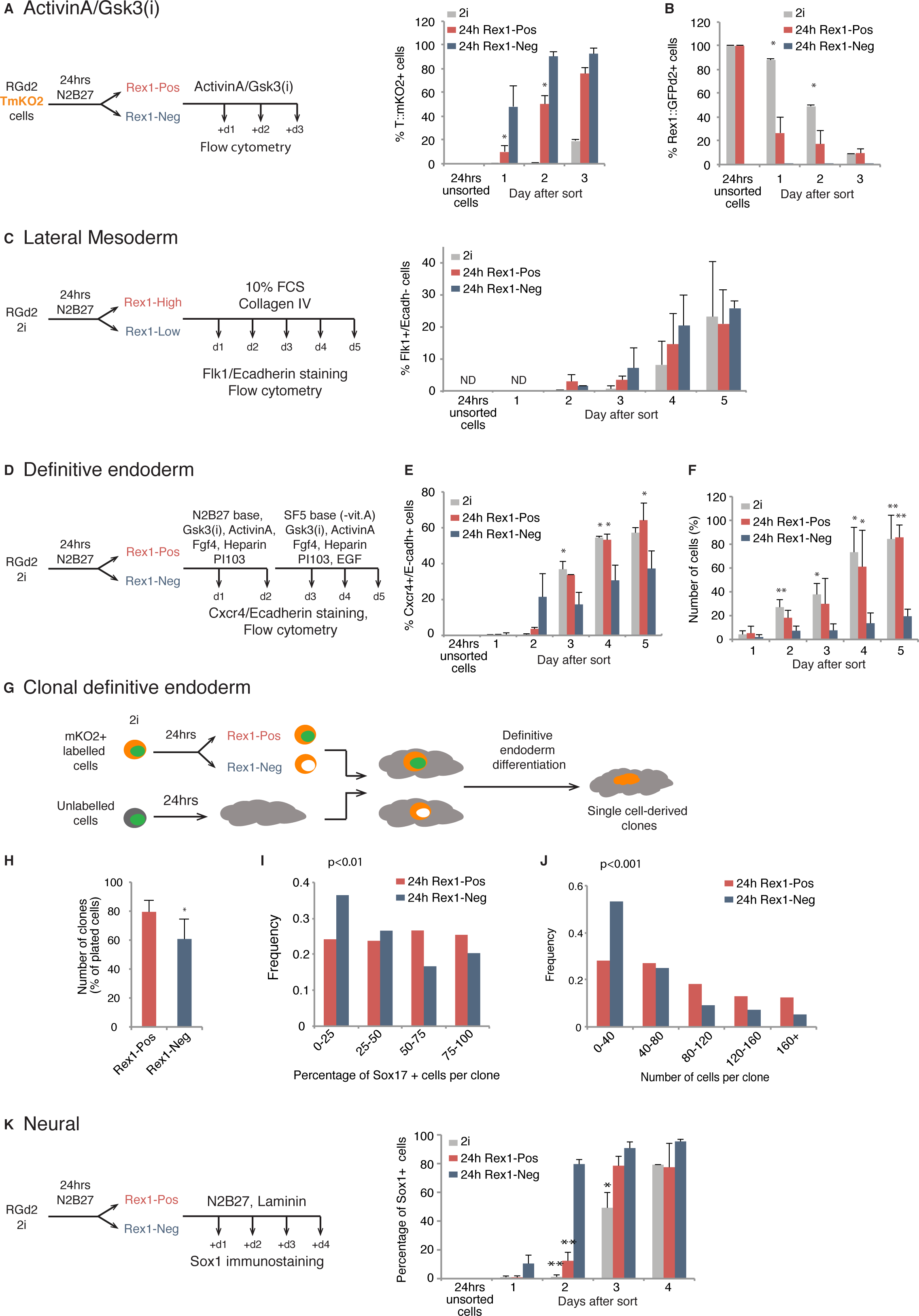
Multilineage differentiation capacity is manifest in Rex1-negative cells. (**A**) Experimental set up and sample analysis for ActivinA+GSK3(i) treatment. Histogram shows the percentages of cells expressing *T:mKO2* or *RGd2* (**B**). (**C**) Experimental set up and sample analysis for lateral mesoderm differentiation. Histogram showing the percentage of Flk1+/Ecadh-cells. (**D**) Experimental set up and sample analysis for definitive endoderm differentiation. (**E**) Percentage of Cxcr4+/Ecadh+ double positive cells. (**F**) Normalised number of cells during definitive endoderm differentiation. The number of cells was normalised to the highest value obtained in that biological replicate. (**G**) Single cell analysis during definitive endoderm differentiation by seeding fluorescently labelled Rex1-High or Rex1-Low cells at clonal density amongst unlabelled cells. (**H**) Number of clones after 4 days of differentiation. (**I**) Histogram showing the distribution of the percentage of Sox17 positive cells per clone. Two independent experiments, all data shown. (**J**) Histogram showing the distribution of the number of cells per clone. Two independent experiments, all data shown. (**K**) Experimental set up and sample analysis for neural differentiation. Histogram showing the percentage of Sox1-positive cells during the differentiation time-course. Unless stated, mean and SD for 3 independent experiments shown, * p<0.05, **p<0.01. See also Figure S2.

Differentiation into definitive endoderm was assessed by monitoring the percentage of Cxcr4/E-cadherin double positive cells (Morrison et al., 2008; Yasunaga et al., 2005) under inductive conditions applied after sorting (Morrison et al., 2015)(Figure 2D). Compared to 2i cells or the Rex1-positive population, a lower proportion of Rex1-negative cells upregulated Cxcr4 (Figure 2E). However, we observed that the majority of Rex1-negative cells died after replating in these conditions (Figure 2F). The survivors could form Sox17/Foxa2 double positive cells, although with lower efficiency than 2i or Rex1-positive cells (Figure S2B). Every Sox17 positive cell was also positive for Foxa2, substantiating endoderm identity (Burtscher and Lickert, 2009). Acquisition of the later marker, Sox17, was specifically reduced in the Rex1-negative cells. We hypothesised that Rex1-negative cells might display impaired survival and differentiation because of a requirement for high cell density and cell-cell contact for the endoderm programme. We therefore combined sorted cells with unsorted populations to reproduce the density of non-manipulated cultures (Figure 2G). To trace the sorted cell progeny we employed *RGd2* cells constitutively labelled with mKO2 under the control of a CAG promoter (Niwa et al., 1991). Two hundred sorted labelled cells were plated together with 5.8×10^3^ parental cells per 3.8cm^2^ dish. Cells were exposed to definitive endoderm differentiation media then fixed and stained for Sox17 at day 4. The total number of mKO2 positive clones was determined, as well as the number of Sox17 positive cells per clone and clone sizes using CellProfiler (Jones et al., 2008). Slightly fewer clones were obtained from Rex1-negative cells (Figure 2H, Student t-test p<0.05) and their distribution was skewed towards smaller colonies (Figure 2I, two-sample Kolmogorov-Smirnov test p<0.001), with more Sox17 negative cells per colony (Figure 2J, two-sample Kolmogorov-Smirnov test p<0.01). These differences were modest however. Importantly, the majority of Rex1-negative cells were able to produce colonies containing Sox17 positive cells.

Finally, we examined cell fate acquisition in N2B27 alone, which is permissive for neural differentiation (Ying et al., 2003). The great majority (≤80%) of cells from both Rex1 fractions became immunopositive for Sox1, an exclusive marker of neurectoderm (Pevny et al., 1998; Zhang et al., 2010) (Figure 2K). However, Rex1-negative cells showed earlier upregulation of Sox1, with most cells becoming Sox1 positive on day 2, a day before the Rex1-positive population (Figure 2K). Cell viability and expansion were not significantly different between the populations (Figure S2C). Rex1-negative cells subsequently also showed accelerated onset of expression of the neuronal marker type III *β*-tubulin (Lee et al., 1990)(Figure S2D).

Overall, these data indicate that after 24hrs of monolayer differentiation guided by autocrine cues, cells in the Rex1-negative population are poised for multilineage specification and respond more rapidly to induction than either ground state ES cells or Rex1-positive cells.

### Nodal does not regulate kinetics of exit from the naïve state

FGF4 is a known autocrine factor that drives ES cell transition upon release from 2i (Betschinger et al., 2013; Kunath et al., 2007; Leeb et al., 2014; Stavridis et al., 2007). A second potential autocrine regulator is Nodal (Fiorenzano et al., 2016; Mullen et al., 2011; Ogawa et al., 2007). Detection of Smad2 phosphorylation indicates that endogenous Nodal/TGFβ signalling is active in ES cells in 2i (Figure S3A). Treatment with the Alk5/4/7 receptor inhibitor A83-01 (Alk(i)) (Tojo et al., 2005) eliminated Smad2 phosphorylation after 30 minutes (Figure S3A). However, culture in Alk(i) did not affect colony forming capacity in 2i/LIF, even after continuous culture for three passages (Figure S3B), confirming that Nodal plays little or no role in maintenance of ground state mouse ES cells.

We examined the contribution of autocrine Nodal signalling in progression from the ES cell state. We analysed changes in gene expression in cells withdrawn from 2i in the continuous presence of Alk(i) and found no difference in the dynamics of downregulation of *Nanog* or *Klf2* mRNA (Figure 3B), nor of Nanog and Klf4 protein (Figure 3C). The rate of decay in ES cell clonogenicity was also unaffected (Figure 3D). We conclude that Nodal signalling does not promote initial exit from the naïve state.

**Figure 3.**
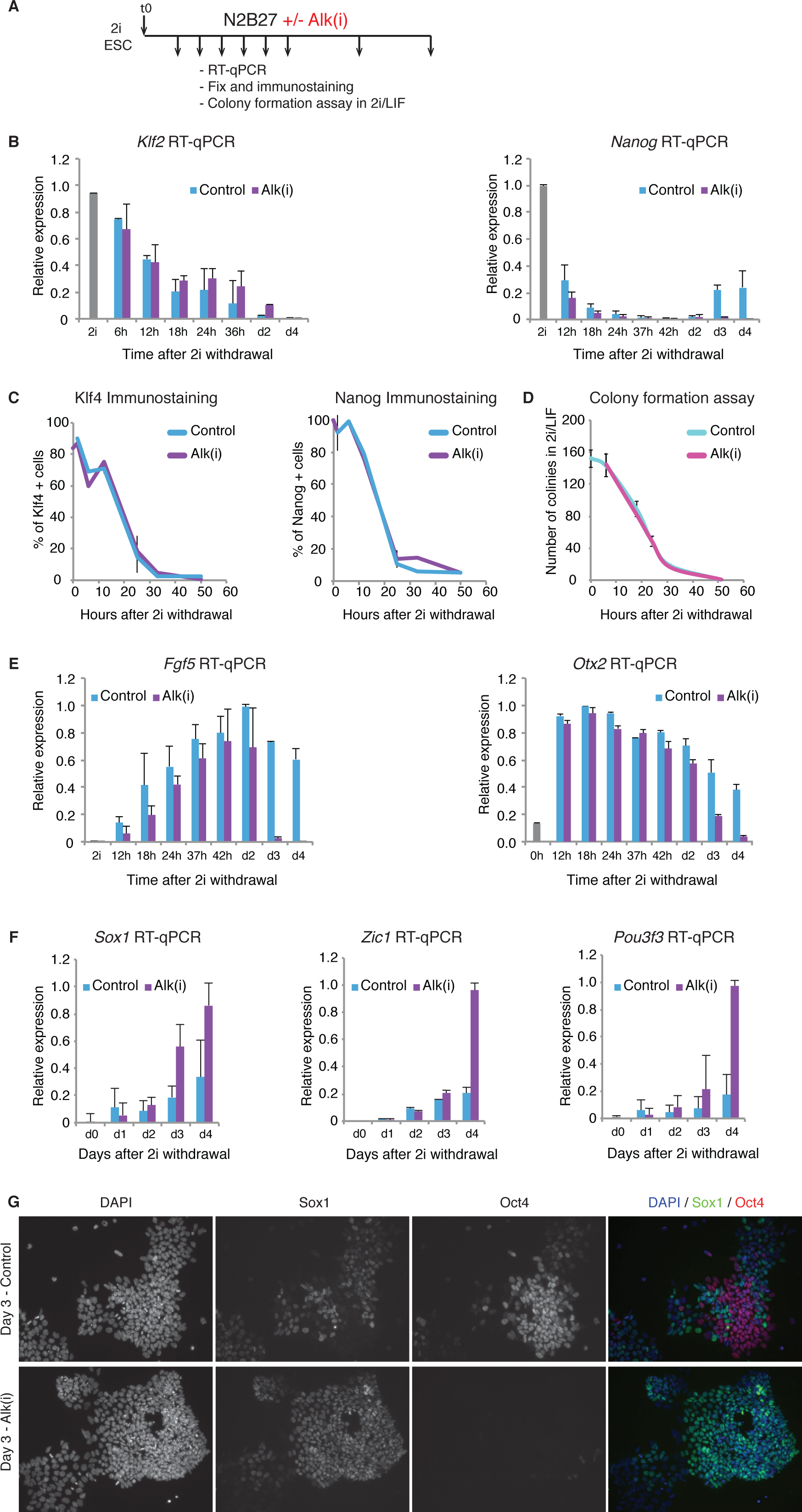
Inhibition of endogenous Nodal signalling does not affect exit from the naïve state. (**A**) Experimental set up (**B**) Relative expression of pluripotency factors *Klf2* and *Nanog* over time when cells are differentiated in DMSO or Alk(i). (**C**) Percentage of Klf4 and Nanog positive cells over time after 2i withdrawal when cells are differentiate in the presence of DMSO or Alk(i). (**D**) Self-renewal capacity declines at a comparable rate for cells treated with DMSO vehicle or Alk(i). (**E**) Relative expression of post-implantation markers *Fgf5* and *Otx2* shows faster earlier downregulated for cells treated with Alk(i) over DMSO controls. (**F**) Relative expression of neural-associated genes *Sox1, Zic1* and *Pou3f3* over time when cells are differentiated in DMSO or Alk(i). (**G**) Inhibition of Nodal signalling results in accelerated reduction of Oct4 protein and increase in Sox1 protein at day 3 of differentiation. Mean and SD for 2 independent experiments shown. See also Figure S3.

We examined expression of genes associated with the early post-implantation epiblast. Initial upregulation of pan-epiblast genes *Fgf5* and *Otx2* was not significantly altered when Nodal signalling was inhibited (Figure 4E). However, these genes were subsequently downregulated more abruptly on day 3/4 (Figure 4E). Conversely, transcripts for neuroectodermal lineage factors *Sox1, Zic1* and *Pou3f3* were strongly up-regulated in day 3/4 Alk(i) treated cultures, before appreciable expression in vehicle treated cells (Figure 3F). At the protein level, most cells in Alk(i) treated cultures had downregulated Oct4 and were Sox1 positive after 3 days, indicative of neural commitment, whereas control cultures displayed a mosaic pattern of co-exclusive Sox1 and Oct4 immunostaining (Lowell, 2006) (Figure 3G).

**Figure 4.**
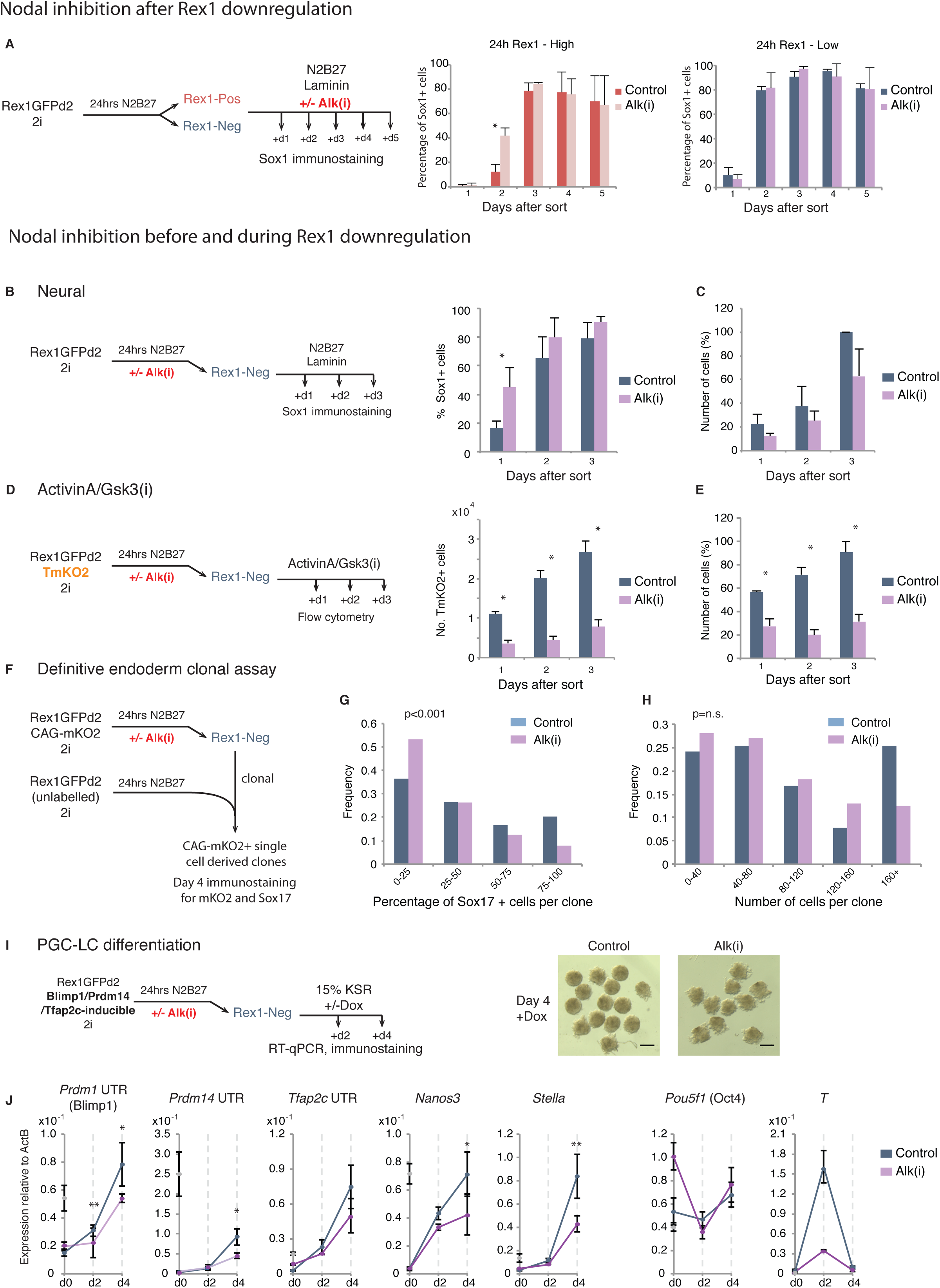
Nodal signalling during exit from the naïve state prevents preconscious neutralisation. (**A**) Inhibition of Nodal signalling with Alki(i) in Rex1-positive and Rex1-negative sorted fractions. Graphs show percentage of Sox1 positive cells after sort. (**B**) Percentage of Sox1 positive cells arising from Rex1-negative cells following DMSO (control) or Alk(i) treatment. (**C**) Number of cells over the period analysed in B. (**D**) ActivinA/Gsk3(i) induction of Alk(i) or control treated Rex1-negative cells. Numbers of TmKO2 positive cells, along with total cell numbers (**E**). To determine the normalised number of cells as a percentage for each biological replicate, the number of cells was normalised by the highest value obtained in that biological replicate. (**F**) Experimental set up of definitive endoderm clonal assay. (**G**) Histogram showing the distribution of the percentage of Sox17 positive cells per clone. Two independent experiments, all data shown. (**H**) Histogram showing the distribution of the number of cells per clone. Two independent experiments, all data shown. Unless states, mean and SD for 3 independent experiments shown, * p<0.05, **p<0.01. (**I**) Experimental set up of transcription-factor dependent PGC-LC differentiation. Images show day 4 cultures in the presence of Dox from Alk(i)-treated and control cells. Scale bar=1mm. (**J**) RT-qPCR of PGC-associated genes during induction process. Mean and SD for 2 independent experiments shown, *p<0.05, p<001. See also Figure S4.

To validate findings with the inhibitor we deployed siRNAs against Nodal signalling pathway components. In *Nodal, Smad2/3* and *Tdgf1* knockdown experiments the emergence of Oct4-/Sox2+ and Sox2+/Sox1+ cells was accelerated (Figure S3C-D). We conclude that suppression of Nodal signalling does not substantially affect initial exit from the naïve state but promotes subsequent specification to the neural lineage.

### Nodal signalling is required to prevent precocious neuralisation and to potentiate other lineages

Examination of Nodal signalling components in RNAseq data from *RGd2* sorted cells (Kalkan et al., 2016) revealed that pathway ligands, receptors, intracellular mediators and target genes are expressed in undifferentiated ES cells and in 24hr Rex1 positive cells. Rex1-negative cells, however, display reduced expression of *Nodal,* Nodal proprotein convertase *Pcsk6* (Pace4), and Nodal signalling pathway targets *Lefty1, Lefty2* and *Smad7* (Figure S4A). Consistent with pathway down-regulation in Rex1-negative cells, we found that when cells were exposed to Alk(i) only after sorting, the Rex1-negative fraction showed no change in kinetics of Sox1 acquisition. In contrast the Rex1-positive population responded by accelerated expression at day 2 (Figure 4A, Student t-test p<0.05).

In light of these results, we postulated that Nodal signalling may function during the primary transition from naïve pluripotency. We therefore inhibited Nodal signalling for only the 24hrs immediately following 2i withdrawal and analysed the resulting Rex1-negative cells (Figure S4B). In line with previous results for continuous treatment, exposure to Alk(i) for 24hrs had little effect on downregulation of Rex1 (Figure S4B) or of naïve pluripotency factor transcripts for *Nanog, Esrrb, Zfp42* (Rex1) and *Klf4* (Figure S4C). Upregulation of early post-implantation markers *Fgf5, Dnmt3b, Otx2* and *Pou3f1* was also similar to vehicle-treated cells (Figure S4C). *Sox1* mRNA was not detected at 24hrs, irrespective of the presence of Alk(i) (Figure S4D). Sox1 protein was detectable only in a minority of untreated cells on day 1 after sorting and increased thereafter. In contrast up to half of cells generated after Alk(i)-treatment upregulated Sox1 protein on day 1 (Figure 4B). This difference does not appear to be due to differential replating efficiency (Figure 4C).

We examined whether faster neural specification as a consequence of Alk(i) pre-treatment has consequences for other lineages. We analysed the response of Alk(i)-treated cells to ActivinA/Gsk3(i). Rex1-negative cells showed a major reduction in the total number of *T::mKO2* positive cells (Figure 4D). Interestingly, this was mainly attributable to reduced cell numbers after exposure to ActivinA/Gsk3(i) (Figure 4E, S4E). A similar reduction in cell survival/proliferation was observed in cells exposed to lateral mesoderm differentiation conditions (Figure S4F-H). To evaluate endodermal specification we employed the clonal mixing protocol described previously (Figure 4F). We observed a shift to fewer Sox17 positive cells per clone (Figure 4G), although the clone sizes (Figure 4H) or total number of clones (Figure S4I) were not reduced in the Alk(i) pre-treated population

Finally, we assessed whether pre-treatment with Alk(i) for 24hrs affected the potential of Rex1-negative cells to respond to PGC-inducing transcription factors (Figure 4I). Alk(i)-treated cells produced less compact and smaller aggregates than control cultures (Figure 4I). The gene expression profile at day 2 and 4 of culture showed lower upregulation of endogenous *Prdm1* (Blimp1), *Prdm14, Nanos3* and *Stella,* indicating significantly impaired PGCLC induction.

These findings indicate that suppression of Nodal signalling reduces the capacity of cells exiting the naïve phase of pluripotency to respond productively to inductive cues for mesoderm, endoderm, and germ cell specification.

## DISCUSSION

The defined context of ground state ES cell culture provides opportunities for experimentally dissecting the interplay between intrinsic and extrinsic factors that mediate progression through pluripotency. Here we investigated the trajectory of ES cells released from the ground state with minimal extrinsic input. We isolated cells that have lost ES cell identity within 24hrs based on down-regulation of *RGd2*, corroborated functionally by extinction of self-renewal capability (Kalkan et al., 2016). Newly formed Rex1-negative cells exhibited capacity for differentiation into the germline and somatic lineages (Smith, *in press*). The findings further indicate that endogenous Nodal signalling is crucial for the non-neural competence of cells transitioning from naïve pluripotency.

Rex1-negative cells show more rapid upregulation of lineage markers in response to inductive stimuli compared with ground state ES cells or Rex1-positive cells at 24hrs. They have also gained capacity for PGCLC induction. It has previously been established that responsiveness to germ cell induction cues or factors is not manifest in naive ES cells or the pre-implantation epiblast but is a property acquired during developmental progression (Hayashi et al., 2011; Nakaki et al., 2013). We present evidence elsewhere that early Rex1-negative cells show intermediate gene expression features suggesting they are related to the peri-implantation epiblast (Kalkan et al., 2016). We hypothesise that actual competence for germline and somatic lineage specification is acquired during this period (Smith, *in press*). The molecular nature of competence remains unclear but is likely to involve dissolution of naïve pluripotency transcription factor circuitry, reconfiguration of the enhancer landscape, and widespread epigenome and chromatin modification (Buecker et al., 2014; Choi et al., 2016; Dunn et al., 2014; Zylicz et al., 2015).

Nodal plays pleiotropic roles in the early embryo. Expression can be detected in the inner cell mass and persists throughout the epiblast until axis specification, when it becomes restricted to the proximal posterior region (Conlon et al., 1994; Mesnard et al., 2006). Nodal activity relies on proprotein convertases, Furin and PACE4, produced by the extraembryonic ectoderm (ExE), which cleave and activate pro-Nodal (Beck et al., 2002; Mesnard et al., 2011). *Nodal* deficient embryos show embryonic lethality at E7.5 (Conlon et al., 1994; 1991; Zhou et al., 1993). They fail to specify the anterior visceral endoderm (AVE) (Brennan et al., 2001), a signalling centre essential for the establishment of anterior-posterior (AP) polarity. Nodal mutants also show precocious upregulation of neural markers throughout the egg cylinder and fail to form a primitive streak (Brennan et al., 2001; Camus et al., 2006; Lu and Robertson, 2004). The multiple functions of Nodal and the complex interplay between extraembryonic tissues and the epiblast have complicated precise delineation of its roles in pluripotency progression and lineage specification (Robertson, 2014).

Mouse ES cells express Nodal and have phosphorylated Smad2/3 proteins (Mullen et al., 2011; Ogawa et al., 2004). Inhibition of Nodal signalling enhances Sox1 expression during differentiation (Matulka et al., 2013; Turner et al., 2014). Our results show that inhibition of endogenous Nodal signalling does not affect the downregulation of pluripotency factors when ground state ES cells are released from 2i, consistent with previous findings (Turner et al. 2014). Upregulation of early post-implantation markers is also unaffected. However, suppression of Nodal signalling results in compromised responses to inductive stimuli for mesoderm and endoderm, and in precocious upregulation of neural markers. Cells also become less responsive to the forced expression of PGC-specific transcription factors.

Importantly, a requirement for Nodal signalling is apparent prior to exit from the ES cell state, while cells are in the reversible Rex1 positive period of transition (Kalkan et al., 2016; Martello and Smith, 2014). Indeed, subsequent to exit Rex1-negative cells in vitro down-regulate Nodal and become dependent on exogenous ligand for non-neural lineage induction, typically achieved by addition of ActivinA. A similar reduction in the expression of *Nodal* and Nodal target genes is seen in E5.75 epiblast explants after removal of the extraembryonic ectoderm (ExE) (Guzman-Ayala et al., 2004; Mesnard et al., 2006), highlighting the paracrine role of ExE in maintaining Nodal signalling in the embryo.

Our findings in the simple monolayer ES cell system are consistent with genetic evidence that Nodal signalling prevents premature neural differentiation in the embryo (Camus et al., 2006). Importantly, however, they also indicate that endogenous Nodal signalling acts during progression from naïve pluripotency to secure non-neural lineage potency. It has been reported that Smad2/3 is recruited by ‘master transcription factors’ to regulatory loci in a cell type-specific manner (Mullen et al., 2011). In addition, a recent study in human ES cells also suggested that Smad2/3 is able to recruit histone methyltransferases to gene promoters (Bertero et al., 2015). Therefore, non-neural competence could depend upon the presence of Smad2/3 at specific loci during the ES cell transition from naïve pluripotency.

Overall these results are consistent with the proposition that in defined adherent culture, ES cells transit through a formative phase in which they acquire competence for multilineage differentiation, including the germline (Kalkan and Smith, 2014; Smith, *in press).* In this phase, cells are expected to respond to inductive signals rapidly and efficiently, as observed for Rex1 negative cells at 24hrs. Furthermore, our findings highlight a pivotal requirement for Nodal signalling in establishing formative pluripotency.

## EXPERIMENTAL PROCEDURES

### Mouse ES cell culture and differentiation

*RGd2* ES cells were derived in 2i/LIF from heterozygous embryos (Kalkan et al., 2016). The *RGd2/T:mKO2* cell line was generated by targeting the endogenous T locus with T2A-mKO2. ES cells were routinely maintained on gelatine-coated plates (Sigma, cat. G1890) in N2B27 media (Stem Cells inc, SCS-SF-NB-02) supplemented with 1μM PD0325901 and 3μM Chir99021 (2i) without LIF, and passaged with Accutase (Millipore, SF006) every 2-3 days. For sorting experiments, cells were plated for 24hrs in 2i at 1.5x10^4^ cells/cm^2^ before washing once with PBS and changing the media to N2B27. After 24-26hrs, cells were sorted by flow cytometry according to GFP levels into Rex1-positive (highest 15%) and Rex1-negative (lowest 15%) populations using a MoFlo sorter (Beckman Coulter, inc). For neural differentiation, cells were plated at 1.0x10^4^ cells/cm^2^ on laminin-coated dishes (Sigma-Aldrich, L2020) in N2B27. Medium was changed every other day. Definitive endoderm induction was as described (Morrison et al., 2015). Lateral mesoderm differentiation was performed by plating 1.2x10^4^ cells/cm^2^ cells in collagen coated plates (BD BioCoat, 354591) in batch tested 10% Serum medium (GMEM (Sigma-Aldrich, G5154), 10% FCS (Sigma-Aldrich), 1x NEAA (Life Technologies, 11140-050), 1mM sodium pyruvate (Life Technologies, 11360-070), 1mM L-Glutamine (Life Technologies, 25030-081)) (Nishikawa et al., 1998).

ActivinA 10ng/ml and Chir99021 3μM (Gsk3(i)) treatment of sorted fractions was carried out on fibronectin-coated plates (Millipore, FC010) at 1.5x10^4^ cells/cm^2^. Nodal inhibitor experiments were carried out using A8-301 1μM (Alk(i), Tocris Bioscience, 2939) or DMSO (1:10000) as a carrier control.

Colony forming assays were conducted by plating 1000 cells per well in laminin-coated 6 well plates in 2i supplemented with 100U/ml LIF to maximise self-renewal potential (Wray et al., 2011). After 5 days, cells were stained using alkaline phosphatase kit (Sigma, cat. 86 R-1KT) and the number of colonies counted.

For transcription factor induction of PGCLC, the tri-cistronic Ap2g-T2A-Prdm14-P2A-Blimp1 fragment (APB1, kind gift from Toshihiro Kobayashi and Azim Surani) was cloned into phCMV^*^1-cHA-IRES-H2BBFP plasmid. pPyCAG-PBase, pPBCAG-rtTA-IN and phCMV^*^1-APB-IRES-H2BBFP were co-transfected into *RGd2* cells by TransIT-LT1 (Kinoshita et al., 2015). G418 selection (400 μg/ml) was started 48 hours after transfection and cells were replated at clonal density at 96 hours. For PGCLC induction, cells sorted at 24 hours for Rex1-GFP expression were plated at 3,000 cells per well in a 96 round-bottomed well plate with (Nakaki et al., 2013) in the presence or absence of 1μg/ml Doxycycline (Sigma-Aldrich). Cells were fed every other day. Aggregates collected on day 2 and 4 for RT-qPCR or fixed after 4 days in culture.

### Flow cytometry analysis of fluorescent reporters

Cells were dissociated into a single cell suspension using Accutase and resuspended in PBS+5% FBS for analysis using a BD LSR Fortessa Analyser.

### Immunohistochemistry

Samples were fixed with 4% PFA for 10min at room temperature (RT), permeabilsed and blocked for 2hrs with block buffer (PBS+0.03%TritonX+3% donkey serum). Cells were incubated overnight at 4 °C in block buffer with the following primary antibodies: Sox1 (Cell Signalling, 4194, 1:200), Oct4 (Santa Cruz, sc-5279 or sc-8628, 1:400), Nanog (eBioscience, 14-5761-80, 1:200), Klf4 (Abcam, ab72543, 1:300), Tuj1 (R&D, MAB1195, 1:500), Foxa2 (Abcam, ab40874, 1:200), Sox17 (R&D, AF1924, 1:200), T (R&D, AF2085, 1:200), Esrrb (Perseus, PP-H6705-00, 1:300), mKO2 (Amalgaam-MBL, M168-3, 1:1000), Blimp1 (eBiosciences, 14-5963-82). After three washes with PBS+0.03%TritonX, cells were incubated with secondary antibodies (Life Technoligies, 1:1000) and DAPI in blocking buffer for 3hrs in the dark. After three washes with PBS+0.03%TritonX, cells were left in PBS before imaging. Images were acquired with Laica DMI3000 B inverted microscope and the fluorescence in single cells quantified using CellProfiler (Jones et al., 2008). The number of cells was normalised to the highest value obtained for a given biological replicate.

### Immunostaining of surface markers for flow cytometry

Cells were dissociated with enzyme-free Cell Dissociation Buffer (Life Technologies, 13151-014) at 37 °C. Cells were resuspended with staining buffer (PBS+1% Rat serum) and incubated with directly conjugated antibodies for 30min at 4 °C in the dark. After three washes with staining buffer, cells were analysed on an LSR Fortessa (BD Bioscineces). Spherotech beads were used to quantify the number of cells. The following antibodies were used: Ecadherin-eFluo660 (eBioscience, 50-3249-82), Cxcr4 (BD Biosciences, 552967 or 558644), Flk1 (BD Biosciences, 562941).

### Gene expression analysis

RNA isolation from cell populations was performed with RNAeasy kit (Qiagen). SuperScriptIII (Invitrogen) and oligo-dT primers were used to synthesise cDNA. TaqMan probes were used for *Pou5f1* (Oct4), *Sox2, Nanog,* Esrrb, *Zfp42* (Rex1), *Klf2*, *Otx2, Fgf5, Pim2, Sox1* and *Dnmt3b.* UPL primers were used for *Pou3f1* (fw: catttttcgtttcgttttaccc, rv:gagcgcagaccctctctg, probe:72), *Smad2* (fw:aggacggttagatgagcttgag, rv: gtccccaaatttcagagcaa, probe:9), *Tdgf1* (fw: gtttgaatttggacccgttg, rv:ggaaggcacaaactggaaag, probe:93), *Nodal* (fw: ccaaccatgcctacatcca, rv:cacagcacgtggaaggaac, probe:40), *Lefty2* (fw: cacaagttggtccgtttcg, rv:ggtacctcggggtcacaat, probe:78), *Zic1* (fw: ggtacctcggggtcacaat, rv:cctcgaactcgcacttgaa, probe:7), *Pou3f3* (fw: tctgagaccgcccacaag, rv: gagcggcagtcagcaaag, probe:22).

### Gene knockdown

Qiagen FlexiTube siRNAs for *Nodal, Tdgf1, Smad2* and *Smad3* at a final concentration of 20nM were used for gene knockdown. 1.5x10^4^ cells/cm^2^ were transfected in 24 well plates containing 500μl of medium 2i medium with 0.5μl Lipfectamine RNAiMAX (Life Technologies, 13778075) for. After overnight incubation, cells were washed once with PBS before transfer to N2B27. Efficiency of transfection was quantified by flow cytometry on Rex1GFPd2 cells transfected overnight with siRNA against GFP. Gene knockdown was quantified by RT-qPCR after overnight transfection.

### Immunoblotting

Western blotting was performed using standard techniques. The following primary antibodies were used: Smad2 (Cell Signalling 3101, 1:1000 in 1% milk), p-Smad2 (Cell Signalling, 3103, 1:1000 in 1% milk), anti-GAPDH (Sigma-Aldrich, G8795, 1:5000 in 1% milk). Peroxidase-conjugated secondary antibodies were used (Sigma-Aldrich, 1:5000). Amersham ECL Western Blotting detection reagent (RPN2106) was used according to manufacturers instructions.

### Statistics

ANOVA was used to compare three or more samples. Two-tailed Student's t test was used for pairwise comparisons. Kolmogorov-Smirnov test was used to determine statistical significance of endoderm differentiation mixing experiments.

## AUTHOR CONTRIBUTIONS

C.M., T.K. and A.S. designed the experiments. C.M. performed the experiments, analysed the data and prepared figures. A.S. supervised the study. C.M. and A.S. wrote the paper.

## ACKNOWLEDGEMENTS

We thank Andy Riddell and Nigel Miller for flow cytometry and FACS support and Peter Humphreys for imaging support. We also thank Martin Leeb for creating and validating the TmKO2 reporter and Masaki Kinoshita for targeting and validated the *Rex1::GFPd2* APB1 cell line. The APB1 construct was a kind gift from Toshihiro Kobayashi and Azim Surani. We thank Jennifer Nichols, Kevin Chalut, Graziano Martello and Harry Leitch for discussions and comments on the manuscript. This research was funded by the Wellcome Trust. The Cambridge Stem Cell Institute receives core support from the Wellcome Trust and the Medical Research Council. C.M. was funded by a BBSRC studentship. A.S. is a Medical Research Council Professor.

## SUPPLEMENTARY FIGURE LEGENDS

**Figure S1**

(**A**) Flow cytometry histogram of RGd2 cells after removal of 2i.

(**B**) Experimental set up for sorting experiments.

(**C**) Flow cytometry profile of sorted fractions

(**D**) Replating capacity of 2i, 24hrs Rex1-positive and 24hrs Rex1-negative cells in 2i/LIF media.

(**E**) Replating capacity of 2i, 24hrs Rex1-positive and 24hrs Rex1-negative cells in Serum/Lif media.

(**F**) Quantification of Oct4 immunostaining in day 4 aggregates in the presence of Dox. Mean and SD for 3 independent experiments shown.

**Figure S2**

(**A**) Flow cytometry plots of Rex1GFPd2+TmKO2 cells in 2i, and Rex1-positive sorted cells for 3 days in control or ActivinA+GSK3(i).

(**B**) Percentage of cells staining positive for Sox17 and Foxa2 during definitive endoderm differentiation.

(**C**) Normalised number of cells during neural differentiation.

(**D**) Immunostaining for Sox1 and Tuj1 of 2i, Rex1-positive and Rex1-negative cells after 6 and 8 days of differentiation.

To determine the normalised number of cells as a percentage for each biological replicate, the number of cells was normalised by the highest value obtained in that biological replicate.

Mean and SD for three independent experiments shown.

**Figure S3**

(**A**) Western blot showing p-Smad2, Smad2 and Gapdh protein in cells treated with control or Alk(i) for 30min.

(**B**) Number of colonies of ES cells grown in 2i+DMSO or 2i+Alk(i) for three passages. Mean and SD for two independent experiments is shown.

(**C**) RT-qPCR of *Nodal, Tdgf1* and *Smad2/3* siRNA treated cells after overnight transfection in 2i. siRNA knockdown did not affect the expression of pluripotency genes *Pou5f1, Klf4* or *Nanog* but in some cases it did affect the expression of the Nodal signalling target *Lefty2.*

(**D**) Quantification of the number of Oct4/Sox2 double positive cells, Oct4 negative/Sox2 positive and Sox2/Sox1 double positive cells on day 3 of neural differentiation after treatment with siRNA. Mean and SD for two independent experiments is shown. * p <0.05, ** p<0.01

**Figure S4**

(**A**) Expression of Nodal pathway signalling components in 2i, Rex1-positive and Rex1-negative cells (Kalkan et al.).

(**B**) Nodal inhibition before and during downregulation of Rex1 – Flow cytometry plot of Rex1GFPd2 cells differentiated for 24hrs in Alk(i) or control (DMSO).

(**C**) Relative expression of pluripotency and differentiation factors in Rex1-negative cells arising from control or Alk(i) conditions by RT-qPCR. 2i and Rex1-positive cells are included as controls.

(**D**) Expression of *Sox1* in the sorted fractions.

(**E**) Percentage of TmKO2 positive cells during ActivinA/Gsk3(i) treatment of Control or Alk(i) derived Rex1-negative cells. Mean and SD for 3 independent experiments shown, *p<0.05.

(**F**) Lateral mesoderm differentiation of 24hrs Alk(i) or control treated Rex1-negaive cells.

(**G**) Percentage of Flk1+/Ecadh-cells.

(**H**) Histogram showing the normalised number of cells.

(**I**) Number of clones after 4 days of definitive endoderm differentiation.

To determine the normalised number of cells as a percentage for each biological replicate, the number of cells was normalised by the highest value obtained in that biological replicate. Mean and SD for 3 independent experiments shown, *p<0.05.

